# Topolow: A mapping algorithm for antigenic cross-reactivity and binding affinity assay results

**DOI:** 10.1101/2025.02.09.637307

**Authors:** Omid Arhami, Pejman Rohani

## Abstract

**Motivation:** Understanding antigenic evolution through cross-reactivity assays is crucial for tracking viral pathogens and informing vaccine development, particularly for rapidly evolving pathogens requiring regular vaccine updates. However, existing cartography methods, commonly based on multidimensional scaling (MDS), face significant challenges with sparse data, producing incomplete and inconsistent maps. There is an urgent need for robust computational methods that can accurately map antigenic relationships from incomplete experimental data while maintaining biological relevance, especially given that up to 95% of possible measurements may be missing in large-scale studies.

**Results:** We present Topolow, a physics-inspired optimization framework that transforms cross-reactivity and binding affinity measurements into accurate positions in a phenotype space. By modeling antigenic relationships as a system of particles connected by springs representing measured similarities, with selective repulsion between unmeasured pairs, Topolow achieves superior performance compared to MDS. Applied to neutralization data for H3N2 influenza and HIV viruses, our method demonstrated 12% and 43% improvement in prediction accuracy respectively, while maintaining complete positioning of all antigens. Topolow determines optimal dimensionality through likelihood-based estimation, avoiding distortions due to insufficient dimensions, and demonstrates orders of magnitude better stability across multiple runs. The method effectively reduces experimental noise and bias, revealing the underlying antigenic relationships and clusters.

**Availability and implementation:** Topolow is implemented in R and freely available at [https://github.com/omid-arhami/topolow]. The package is optimized for both single machines and SLURM clusters, with parallel processing support for computational efficiency.

**Supplementary information:** Supplementary data are available.

## 1 Introduction

Understanding and tracking the antigenic evolution of viruses is crucial for public health, particularly for rapidly evolving pathogens like influenza that require regular vaccine updates [1, 2, 3, 4]. The ability of viruses to escape immune recognition through antigenic change enables them to reinfect previously exposed individuals [5, 6]. This process, known as antigenic drift, is quantified through cross-reactivity assays that measure how well antibodies generated against one virus isolate recognize and neutralize other strains [7, 8, 9].

The outcome of these assays is usually expressed as either titers or concentrations. A titer refers to the highest dilution of a sample (e.g., serum or antibody solution) that still produces a measurable effect, such as neutralizing viral activity or inhibiting hemagglutination. In contrast, a concentration such as *IC*_50_ (half-maximal inhibitory concentration) is a quantitative measure of the concentration of a substance required to inhibit a biological function by 50%.

Given the resource-intensive nature and substantial costs of these assays, only a small fraction of possible pairwise antigenic relationships are measured directly. Assay measurements also exhibit significant experimental variability [10, 11]. This set of non-complete and noisy pairwise immunological measurements forms a complex network of relationships that can be difficult to interpret directly.

The development of antigenic cartography, primarily through multidimensional scaling (MDS), has been instrumental in visualizing and analyzing antigenic relationships among viruses [1]. MDS-based methods project immunological measurements into a continuous low-dimensional space where antigenic distances between viruses correspond to their immunological differences. A compelling demonstration of the MDS approach to antigenic cartography was provided by Smith et al. [1], who used gradient descent to minimize the sum of squared errors between predicted and measured distances. The method does, however, face challenges with sparse data [12] - a common issue as dataset size increases [13].

Data sparsity arises since assays such as Hemagglutination Inhibition (HI) are typically constrained to a limited number of contemporary antigens. When separate HI tables are merged into a super-matrix, the resulting table is generally highly incomplete, with up to 95% missing values in datasets spanning multiple decades [13]. The abundance of missing values in a dataset forms one of the most significant barriers to creating accurate antigenic maps [12, 13].

Thinking of cartography from a different perspective helps understand the problem of missing measurements. Creating a map is equivalent to determining the coordinates of the points in an *r*-dimensional space, where *r* can be any integer greater than 1. If we have *c* points and their similarities to, or distances from, *r* references are fully measured, the coordinates of points can be exactly determined in an *r*-dimensional space. However, if only a part of similarities/distances between points and references are available, each point can assume an infinite number of positions in the *r*-dimensional space, and there will not be a unique solution. In this case, the common approach is to use MDS to find the positions in a lower-dimension space [14]. However, missing data creates a challenging trade-off for MDS between accuracy and completeness in dimensionality selection: Choosing more dimensions for the target space increases accuracy but hinders finding positions for all points when the number of measurements is smaller than the number of dimensions. Conversely, choosing lower dimensionality causes loss of information and compromises the accuracy of the estimated positions. As the proportion of missing data increases, the dimensionality selection becomes more challenging due to insufficient constraints from the observed distances. Further, as a gradient-based method, MDS is also adversely affected by missing data, which impact the accuracy of the magnitude and directions of gradient vectors.

Additionally, as we demonstrate in this paper, antigenic maps created by MDS for the same data vary substantially between runs. This convergence instability, combined with relatively high mapping errors, affects our ability to visualize accurately and understand antigenic evolution with confidence, ultimately impacting critical public health decisions in vaccine development and viral surveillance efforts.

Several variants have been developed to improve performance relative to MDS in the context of antigenic data, such as use of non-metric MDS [14], temporal matrix completion [13], Bayesian MDS [15], biological matrix completion [16], and integrating protein structure [17].

Here, we adopt a novel, physics-inspired optimization framework that transforms cross-reactivity titers and binding affinity values into spatial representations in the optimal dimensionality. Our method is called Topological Optimization for Low-Dimensional Mapping or Topolow. The algorithm estimates the antigenic map through sequential optimization of pairwise distances, reducing the multidimensional problem to a series of one-dimensional calculations. This gradient-free approach eliminates the need for complex gradient computations required by MDS, enabling robust performance even with substantially incomplete assay data. As demonstrated in our results, Topolow achieves superior accuracy compared to MDS when handling missing data.

## 2 Materials and Methods

### 2.1 Data description and preparation

Pairwise dissimilarity measurements, such as HI titers or neutralization *IC*_50_ values, are typically represented in matrix form. The matrix contains three types of entries: (a) numeric values, (b) threshold values that may represent either low (≤ *h*) or high reactors (*> h*), and (c) missing values.

A titer *T*_*ij*_ in HI assay measures the similarity between test antigen *i* and reference antigen *j*; this is transformed into a dissimilarity measure denoted by *D*_*ij*_:

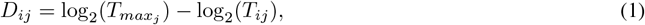

where *T*_*max*_*j* represents the maximum titer value observed for the reference antigen [1]. Since experimental conditions and antiserum potency can vary between assay panels, we search the entire dataset for the maximum titer of each reference antigen rather than using either the homologous titer [2] or the maximum titer within a single panel or clade. This approach helps normalize measurements across different experimental batches.

Threshold values (*h*) are incorporated in the algorithm as equality constraints. Missing values can be predicted by the model once antigenic coordinates are determined.

### 2.2 Proposed mathematical model

The algorithm models antigenic relationships as a physical system where test and reference antigens are represented as similar particles in an *N*-dimensional space, resembling force-directed graph layout approaches [18, 19]. Each pair of particles for which we have a measurement is connected by a spring with a free length equal to their measured dissimilarity, *D*_*ij*_. Following Hooke’s law, the magnitude of the force exerted by the spring is proportional to the displacement from its free length: **F**_*s*,*ij*_ = *k*(*r*_*ij*_ − *D*_*ij*_), where *k* is the spring constant and *r*_*ij*_ is the current distance between particles *i* and *j*.

Pairs of particles lacking a direct measurement apply a repulsive forces to each other that follows the inverse square law: 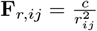, where *c* is a repulsion constant that is fitted from data. This design choice is both biologically sensible and computationally efficient. Previous studies have shown that antigenic distances between temporally distant strains tend to be large [1, 13], yet such pairs are rarely measured in laboratory assays due to logistical constraints. By applying repulsion specifically between pairs with no measurement, our model naturally facilitates separation while avoiding unnecessary force calculations.

For each particle *i*, the total force **F**_*i*_ is the sum of spring forces from its measured neighbors (*N*_*i*_) and repulsive forces from non-connected particles:

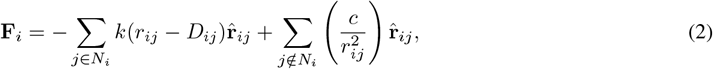

where 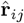 is the unit vector from *i* to *j*.

We assign weights to particles in their motion under the forces, analogous to vertex-weighting schemes used in force-directed graph layouts [20]. Each antigen receives an effective mass *m*_*i*_ proportional to its number of measurements, in contrast with traditional MDS which weights all coordinates equally. This weighting scheme provides a natural regularization, preventing over-fitting; well-measured antigens remain more stable while allowing antigens with fewer measurements greater freedom of movement. The motion of a particle during a time step follows Newton’s second law of motion: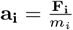, where **a**_**i**_ denotes the acceleration of particle *i*.

A critical distinction between Topolow and force-directed graph layouts is in how distances are treated. In graph layouts, edges typically have uniform lengths and serve mainly to keep connected nodes close together [20]. In contrast, our model implements the true distances through the free lengths of springs, making it suitable for assessing quantitative relationships.

### 2.3 Optimization approach

The algorithm starts by setting initial positions for nodes and optimizes the coordinates to minimize the mean absolute error (MAE) between distances and observations. To initialize particle positions, we leverage existing knowledge about antigenic evolution, which occurs through a combination of genetic drift [15] and directional selection pressure [21].

Rather than a uniformly random initialization, we employ a time-homogeneous Brownian-like Ornstein–Uhlenbeck diffusion process [22] to specify the initial antigenic trait distribution. The initialization affects only the starting positions and does not constrain the final solution, as particles move freely during the optimization process.

Optimization proceeds in the *N*-dimensional space by simulating the physical system’s dynamics. A random permutation of list of particles is created at the beginning of each cycle of simulations. At each time step, *t*, a single particle *i* is selected and its position is updated based on the pairwise interaction forces with one other particle, while all other particles remain fixed. We allow the forces to act on the particle and move it, starting from a stationary state, for one unit of time. Spring and repulsive forces and their induced displacements (*d*_*s*,*i*,*t*_ and *d*_*r*,*i*,*t*_) are calculated as follows:

For spring forces, assuming a constant force during a small displacement results in constant acceleration that can be calculated using Newton’s second law: 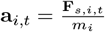. The displacement of the particle during the time step, *d*_*s*,*i*,*t*_, can be derived through the following independent relationships. First, the speed at the end of the time period is calculated as:

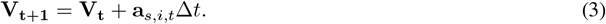

Setting Δ*t* = 1 and initial velocity *V*_*t*_ = 0 yields:

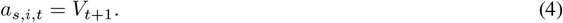

The average velocity in a motion with constant acceleration is the average of the velocities at the beginning and end of the motion:

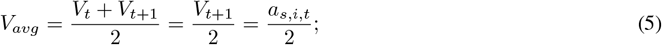

Eq. (4) was used at the last step. We can estimate *d*_*s*,*i*,*t*_ using trapezium rule in discrete numerical integration as *d*_*s*,*i*,*t*_ = *V*_*avg*_Δ*t*. Setting Δ*t* = 1 and using (5) and (4) (in this order):

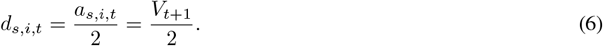

To express *d*_*s*,*i*,*t*_ in terms of the latest distance of particles *i* and *j, r*_*ij*,*t*_, and the constants of the model, we use the rule of conservation of energy:

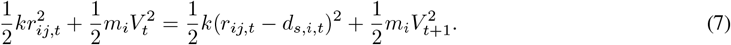

Substituting *V*_*t*_ = 0 and using (4) and (6), gives:

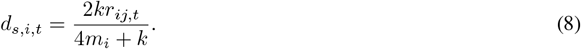

For repulsive forces, Plugging *F*_*r*,*ij*_ in Newton’s second law:

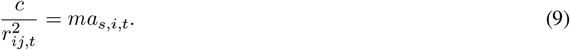

*F*_*r*,*ij*_ is inversely proportional to 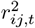, hence, it is comparably small. Consecutively, the resulting displacement 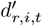 is small compared to *r*_*ij*,*t*_, and we can approximate by assuming a constant force during the motion. Then, similar to (6), we will have 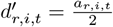. Plugging it in (9)

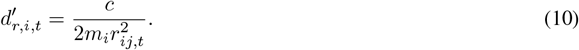

To handle threshold constraints, spring forces are only activated when the current distance falls below the distance corresponding to the threshold value. The force works to push the distance closer to at least the threshold value. (A similar scenario happens in the opposite direction for lower bound thresholds.)

The loss function in Topolow is based on MAE:

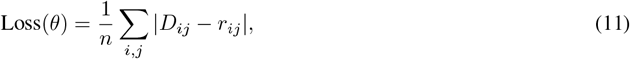

where *θ* represents the model parameters. At the end of each cycle (when the position of all particles were updated), the loss is calculated and the convergence is checked against a threshold. The full pseudo-code of the algorithm and some implementation notes are provided in the supplementary information.

### 2.4 Parameter fitting and Likelihood analysis

There are four parameters in the model: the spring constant (*k*), the repulsion constant (*c*), the dimensionality of the antigenic space (*N*), and the cooling factor (*α*). See “Algorithm and implementation notes” in supplementary information for more detail. An Adaptive Monte Carlo (AMC) sampling approach [23] was employed to construct the likelihood surface for each dataset and determine the optimal value of each parameter, prior to using the algorithm for mapping. Full details are provided in the Supplementary.

Notably, determining the optimal dimensionality, through likelihood analysis is an advancement compared to the convention of choosing 2 or 3 dimensions. Insufficient dimensionality introduces non-uniform distortions in pairwise distances, where some distances become artificially inflated while others are compressed (Fig. S-1). These distortions can generate spurious clusters or merge distinct antigenic groups, potentially confounding biological interpretation.

### 2.5 Simulation study design

To benchmark the relative performance of Topolow and MDS, we carried out a simulation study. Three essential features of antigenic evolution were incorporated in simulated data: directional selective pressure, clustered antigenic phenotypes, and measurement noise [15, 21]. Arbitrary values were selected to generate these features. Dataset complexity was characterized by dimensionality, with antigenic coordinates generated in 2-, 5-, and 10-D spaces. A three-step process was implemented for coordinate generation. Initially, a trend vector was established to represent directional selective pressure and antigenic drift observed in viral data, with linear progression from −10 to 10 arbitrary units across the simulated antigenic space. Five distinct antigenic clusters were then positioned along this trend vector, with cluster centers drawn from a uniform random distribution over the trend length. Biological variation was subsequently introduced through two layers: local spread of antigens around cluster centers was drawn from a multidimensional normal distribution (*σ*_*c*_ = 1), and global phenotypic randomness was implemented (*σ*_*g*_ = 3.3).

For each dimensionality, 250 antigenic phenotypes were generated and split into 150 test and 100 reference antigens to mirror typical experimental conditions. For each dimensional scenario, three datasets with increasing proportions of missing measurements were created (70%, 85%, and 95%). Missing values followed a distance-dependent pattern, preferentially occurring between temporally and antigenically distant antigens. Then three variants for each scenario were developed: (1) “Original” - the base dataset; (2) “+Noise” - original data with added Gaussian noise (*µ* = 0, *σ* = 5% of mean distances); and (3) “+Noise+Bias” - original data with both random noise and a constant bias (5% of mean distances). Table 1 shows a summary of the simulated datasets.

**Table 1:** Parameters and characteristics of simulated datasets.

| Parameter | Value | Description |
| --- | --- | --- |
| Total antigens | 250 | Split into 150 test and 100 reference antigens |
| Dimensions | 2, 5, 10 | Representing different complexity levels |
| Mean antigenic distances | 20.86, 32.60, 46.79 | For datasets in dimensions 2, 5, and 10, respectively |
| SD of antigenic distances | 14.16, 19.18, 27.52 | For datasets in dimensions 2, 5, and 10, respectively |
| Missing data proportion | 70%, 85%, 95% | Percentage of unmeasured relationships |

### 2.6 Cross-validation experimental setup

Model performance was evaluated through 20-fold cross-validation on simulated data. For each of the 27 simulation scenarios, the antigenic distance matrix was randomly partitioned into training (95%) and test (5%) sets. Model parameters were fitted to the training data using AMC simulation (see Supplementary), after which both Topolow and MDS models were employed to predict held-out measurements. Prediction accuracy was quantified using Mean Absolute Error (MAE) for within-data comparisons and Mean Absolute Percentage Error (MAPE) for cross-data evaluations between predicted and true antigenic distances.

### 2.7 Implementation

Topolow was implemented in R 4.3.2 and is freely available as an R package with the same name available at [https://github.com/omid-arhami/topolow]. Topolow is optimized to automatically detect the environment, submit parallel jobs on a SLURM cluster, and use multiple cores on an individual multi-core computer.

As an example of computational resource requirement, fitting model parameters for the H3N2 HI dataset in the Results section using 100 AMC samples, took 80 minutes on a Linux system with 100 CPU cores (2GHz) and 50GB RAM. Map generation was accomplished in 2 seconds on the same machine. Processing time can be reduced further through distributed computing.

## 3 Results

### 3.1 Validation on simulated data

To rigorously benchmark the performance of Topolow, we first designed a comprehensive simulation framework (see Materials and Methods; Fig. S-2).

#### 3.1.1 Performance comparison with MDS

Topolow was evaluated against the commonly used MDS method for antigenic mapping, as implemented in RACMACS [24]. Among existing approaches (e.g., [15, 25]) Topolow and traditional MDS were distinguished by their function as standalone tools for antigenic characterization without requiring additional data types. MDS dimensionality was restricted to 2 dimensions, reflecting both standard implementation and the highest dimensionality at which MDS can position at least 95% of antigens in different datasets, as higher dimensions result in positioning failures for an increasingly large fraction of antigens. As demonstrated in Fig. 2, Topolow consistently achieved significantly lower MAPE than MDS –typically multiple orders of magnitude smaller– across all scenarios (paired t-tests calculated in Tables S-3 and S-4, *p <* 0.0001 for all scenarios). Both methods exhibited decreased accuracy with increasing data complexity (dimensionality) and sparsity. Visual inspection of maps created by Topolow and MDS also shows that while MDS produces distorted relationships and fails to position all points, Topolow maintains the clear cluster structure almost identical to the original data (Fig. S-3 depicts the comparison for the most challenging scenario.)

**Figure 1:**
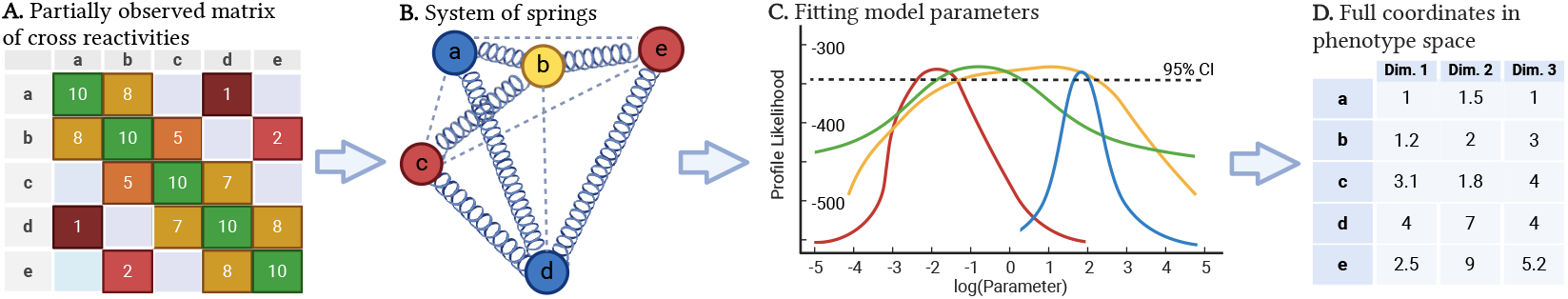
Schematic of the model. (**A**) Assay results are organized into a dissimilarity matrix. (**B**) The matrix is converted into a spring system, in which, antigens are represented as particles connected by springs wherever an assay measurement exists. Dashed lines are missing measurements to be estimated by Topolow. (**C**) The parameters of the model are fitted to maximize the likelihood of the data. (**D**) Topolow finds the optimal dimensionality and coordinates of all test and reference antigens.

**Figure 2:**
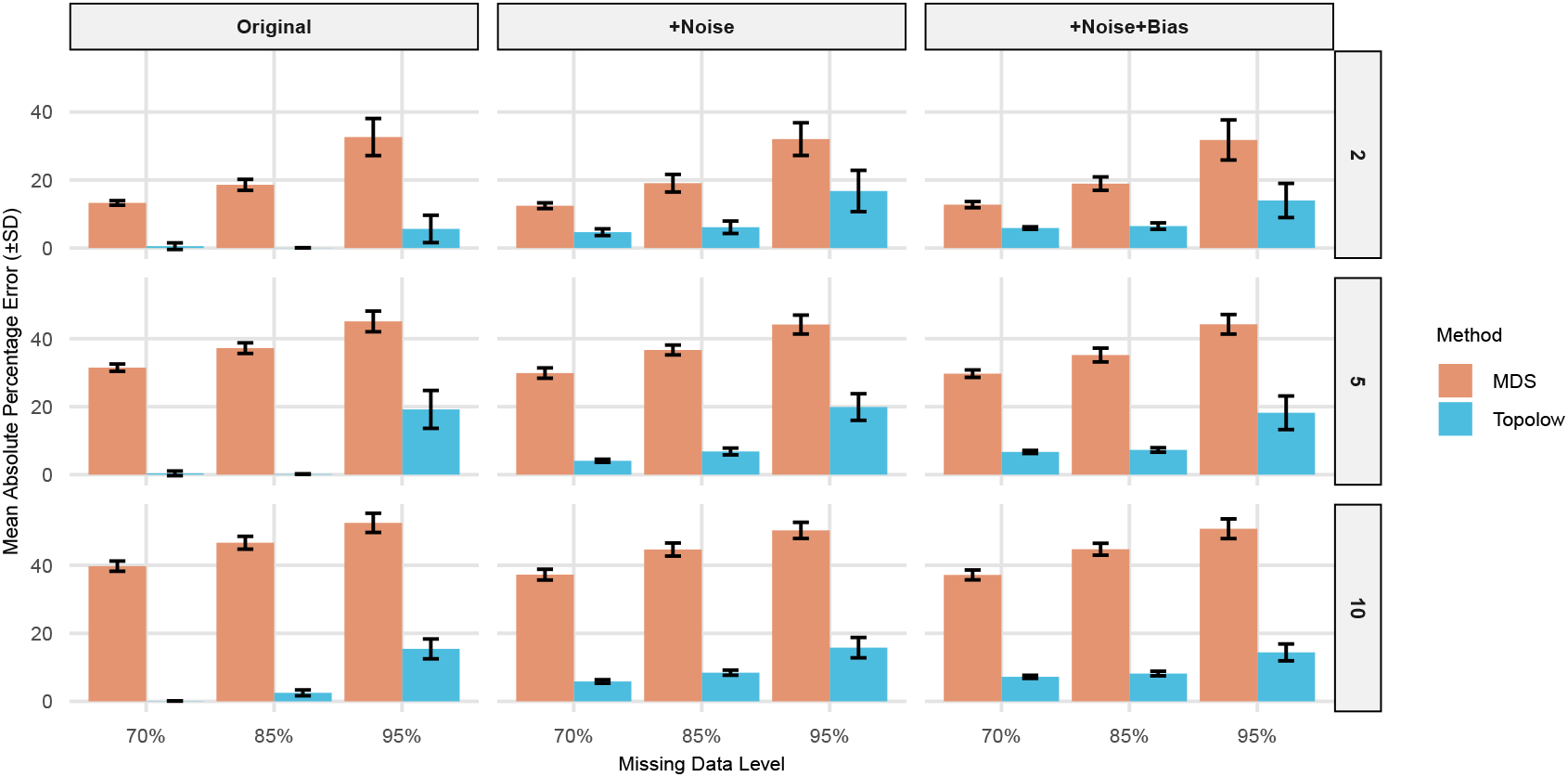
Quantification of model performance on simulated data. MAPE was compared across dimensions 2, 5, and 10, three variants of data: original, +Noise (original distances plus a 5% random noise), +Noise+Bias (original distances plus a 5% random noise and a 5% bias), and three missing percentages: 70%, 85%, and 95%.

#### 3.1.2 Completeness of antigenic maps

A critical feature of Topolow is its complete positioning of antigens regardless of dimensionality or missing data. In contrast, MDS failed to determine positions for an increasing fraction of antigens (up to 99%) in data with greater complexity and missingness, both in 2D and when true data dimensionality was assumed known and used (Table S-1).

#### 3.1.3 Bias analysis

The distributions of error for MDS maps exhibited positive bias across all scenarios, while Topolow consistently achieved near-zero biases (Table S-6). It is notable because antigenic distances are usually compared against a threshold and biases can shift antigens from non-novel to novel area, or vice versa.

#### 3.1.4 Robustness to input noise and bias

One of the potential advantages of creating an antigenic map is the reduction of errors by using multiple measurements to determine the position of each antigen. We test this hypothesis for each method by comparing the MAE of its results against known input errors. Error metrics are defined as follows: (1) Input MAE: The average of absolute differences between distances in the noisy/biased and the original variants of each scenario. This metric represents the baseline noise in the raw data. (2) MDS MAE: The MAE between distances on maps created by MDS and the original distances. (3) Topolow MAE: The MAE between distances on maps created by Topolow and the original distances.

Table 2 shows the comparison of MAEs for 5D scenarios (see other scenarios in Table S-2.) Topolow consistently achieves lower MAE than both the input data and MDS-derived distances, indicating its superior noise reduction capability. This substantial improvement in accuracy demonstrates Topolow’s robust ability to reduce measurement noise in the data and reveal the underlying antigenic relationships.

**Table 2:** Comparison of MAPE (%) in the inputs due to added noise/biase against MAPE of locations determined by MDS and Topolow. Only dimension 5 is shown.

| Dim. | Missing | Variant | Input | MDS | Topolow |
| --- | --- | --- | --- | --- | --- |
| 5 | 70% | +Noise | 5.693 | 29.917 | 3.205 |
| 5 | 70% | +Noise+Bias | 8.488 | 29.769 | 6.507 |
| 5 | 85% | +Noise | 5.420 | 37.041 | 4.073 |
| 5 | 85% | +Noise+Bias | 7.867 | 35.523 | 6.328 |
| 5 | 95% | +Noise | 4.791 | 48.159 | 5.459 |
| 5 | 95% | +Noise+Bias | 7.173 | 47.703 | 7.324 |

### 3.2 Application to Empirical Datasets

We evaluated Topolow using two distinct empirical datasets that represent different challenges in antigenic cartography. The first is the extensively studied dataset of HI assays of H3N2 Influenza antigens from Smith et al. [1], which serves as the gold standard due to its careful curation and validation. The second is a larger, uncurated, HIV-1 subtypes B and C neutralization dataset from Los Alamos National Labs [26], which presents additional challenges of potential noise and the absence of clear antigenic patterns.

#### 3.2.1 H3N2 influenza analysis (1968-2003)

The H3N2 HI dataset [1] contains 4,215 measurements between 273 antigens (test) and 79 antisera (reference), spanning 1968-2003. This represents approximately 23% of all possible test-reference combinations.

Topolow identified these data to be 4D, while two dimensions was used in MDS, as in Smith et al. [1]. Topolow achieved a validation MAE of 0.828 *±* 0.08 *log*_2_ units, comparable with MDS (MAE 0.939 *±*0.11) while yielding a 12% improvement. We note that MDS can reach an MAE=0.85 in 5 dimensions, but it fails to find the location of all antigens on the map (Table S-1). Fig. 3 shows the distribution of errors and detailed performance metrics across methods.

**Figure 3:**
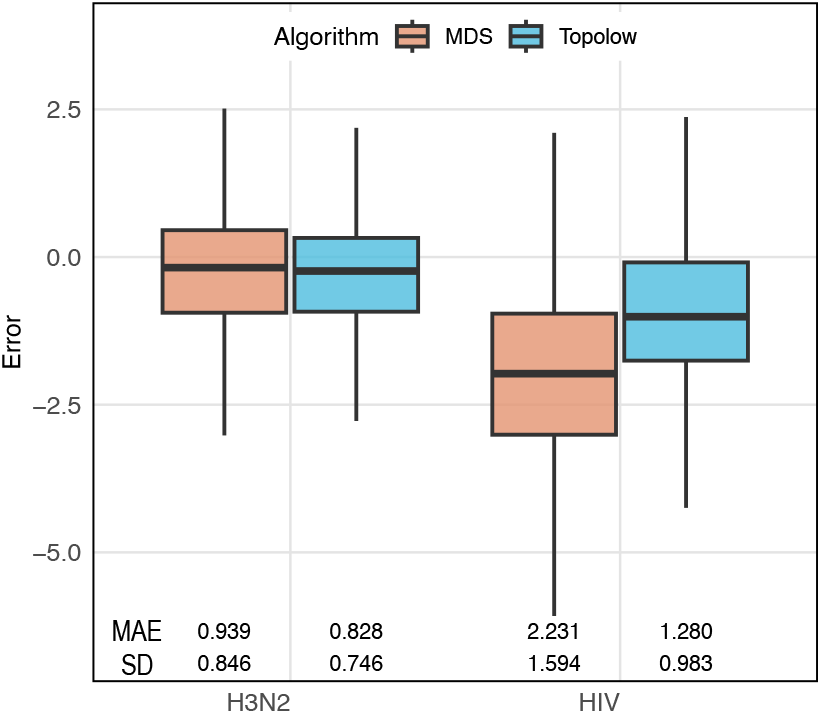
The distribution, MAE, and SD of validation errors across methods, for antigenic distances of H3N2 viruses with 5th to 95th percentile (0 - 7.665) and *log*(*IC*_50_) values for HIV viruses with 5th to 95th percentile (0 - 3.864).

The 2D projection of the antigenic map generated by Topolow is strikingly similar to the map published by Smith et al. [1], using MDS (Fig. 4), with consistently identified key antigenic clusters.

**Figure 4:**
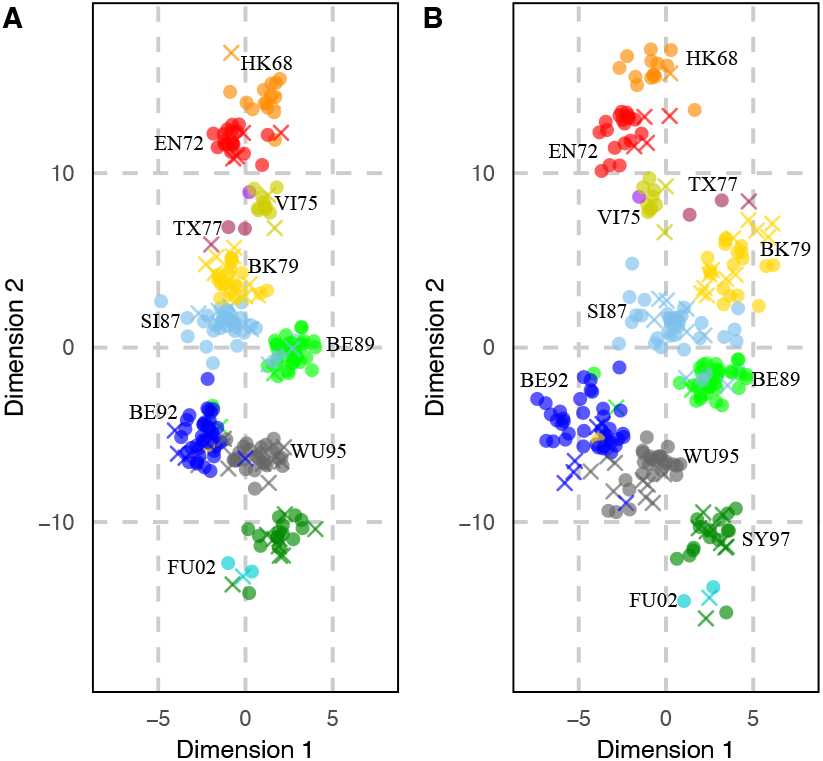
Comparison of antigenic maps estimated by Topolow (**A**) and MDS (**B**) for H3N2 HI titers from 1968 to 2003. Test antigens are shown as colored circles and reference antigens are shown as colored crosses, with colors denoting antigens clusters inferred by Smith et al. [1].

#### 3.2.2 HIV neutralization data

The HIV neutralization dataset comprises *IC*_50_ measurements between 284 antigens and 51 antibodies from HIV-1 subtypes B and C, the subtypes accounting for over 50% of global HIV infections [27]. Only 6.5% of possible test-reference combinations were measured (7,279 measurements total), making this a significantly sparser dataset than that for H3N2.

*IC*_50_ values directly measure antigenic dissimilarity but showed strong right-skew, necessitating log-transformation during pre-processing. Topolow achieved a validation MAE of 1.28 ± 0.983 *log*_2_ units, compared to 2.231 *±*1.594 for MDS - a 43% improvement in prediction accuracy.

The resulting antigenic map (Fig. 5) reveals a pattern of antigenic clustering by viral subtype. Additional analyses (Fig. S-6) expose the clusters more clearly and also demonstrate a temporal trend with increasing antigenic heterogeneity in recent isolates. More detailed analyses of antigenic patterns in Topolow results are warranted but beyond the scope of this paper.

**Figure 5:**
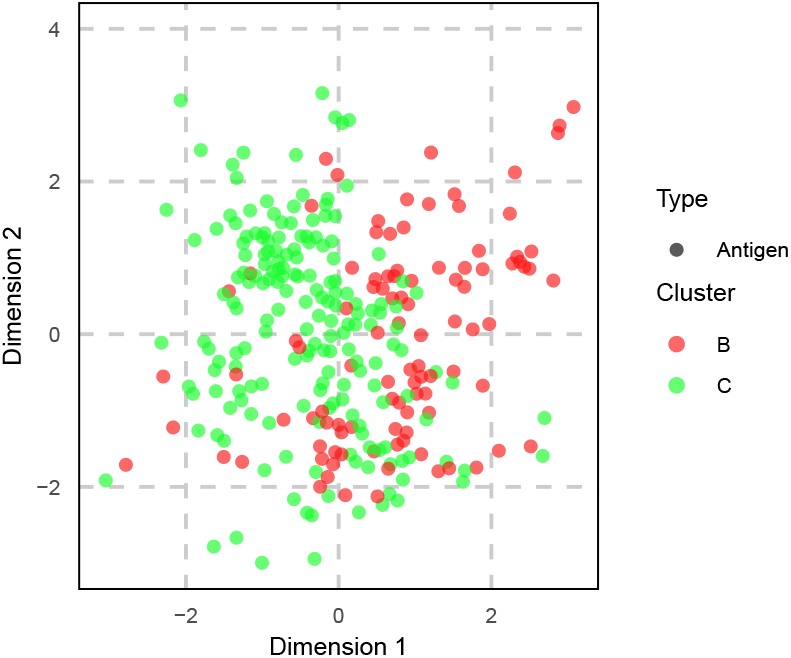
2D visualization of HIV antigenic map created by Topolow, colored by subtypes.

#### 3.2.3 Convergence stability analysis

It is critical for policy applications that antigenic characterizations be consistent across multiple runs of any method. To quantify the stability of Topolow and MDS, 15 independent 2D maps were created by each method for both H3N2 and HIV datasets to have 105 paired maps 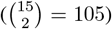for each of the 4 method-data combinations, yielding sufficient power in paired t-tests. MDS was allowed to repeat each run 1000 times and keep the result with the smallest sum of squared errors to avoid convergence to a local optimum. Run-to-run variation for each method-data combination was evaluated by calculating Procrustes sum of squared errors (PSSE) [28] for the 105 paired maps. Since each pair of maps is generated by the same method on the same data, they should not, in principle, differ significantly.

The mean and variance of PSSEs are shown in Table 3. Topolow demonstrated better stability, with mean PSSE multiple orders of magnitude smaller than MDS for both datasets, confirmed by paired t-tests (*p <* 1*e* − 4). The standard deviations (SD) of PSSEs were also substantially lower for Topolow, indicating more consistent performance across runs, confirmed by F-tests for variances (*p <* 1*e* − 4).

**Table 3:** Run-to-run stability analysis (Procrustes sum of squares)

| Data | Method | Mean | SD | t-test |
| --- | --- | --- | --- | --- |
| H3N2 | TopoLow | 9.5 | 7.19 | $p < 1e - 4$ |
| H3N2 | MDS | 3668.56 | 3038.51 |  |
| HIV | TopoLow | 11.89 | 1.21 | $p < 1e - 4$ |
| HIV | MDS | 184.64 | 42.18 |  |

## 4 Discussion

Understanding and quantifying the antigenic evolution of viral pathogens is crucial for public health surveillance and the design of effective intervention strategies [29]. Many rapidly evolving viral pathogens, including influenza [1], SARS-CoV-2 [30], HIV [31], dengue [32], and Hepatitis C [33] require systematic antigenic characterization to guide vaccine development and predict emerging variants while their experimental datasets continue to grow in size, complexity, and incompleteness [34, 31].

Antigenic cartography, often based on MDS, has become a popular tool for interpreting antigenic assay data and producing actionable findings. However, MDS-based approaches are subject to limitations in accuracy, stability, and completeness when applied to large-scale datasets with a high proportion of missing data [13].

Here, we have presented a novel antigenic mapping method, that differs fundamentally from MDS and force-directed graph paradigms. Our method treats antigenic relationships as a physical system of particles connected by springs that move under laws of motion, where the spring lengths represent observed antigenic distances. This topology-based framework optimizes pairwise relationships independently, making it inherently robust to the high proportions of missingness.

Topolow maps uncover the distances between all antigens, including those lacking direct measurements, effectively expanding the usable dataset. This multiplies the training data available for large machine-learning models aimed at understanding immunological relationships and predicting viral evolution, such as [33, 35, 36]. Downstream models with enhanced predictive capabilities combined with more accurate and reliable antigenic maps have the potential to transform how we monitor emerging viral variants and optimize vaccine composition, leading to more effective public health responses to evolving viral threats [37].

Given that the robust detection of clusters in antigenic evolution requires identification of the appropriate dimensionality [38, 39], we propose a key innovation of Topolow is finding the optimal space dimensionality. By approaching the problem as a series of independent one-dimensional simulations –along the line connecting a pair of particles– Topolow circumvents the curse of dimensionality that plagues traditional methods [40]. Unlike MDS, which requires calculating complex gradient vectors in high-dimensional spaces, Topolow only needs to consider movement along the direction connecting two nodes at each time, which allows it to maintain accuracy regardless of the overall dimensionality.

Furthermore, Topolow offers improved results stability compared to MDS which can potentially get trapped in local optima [41], producing different maps for the same data. The mean and variance of Procrustes errors of Topolow were orders of magnitude smaller than MDS. This observation was consistent across both empirical scenarios –the heavily sparse HIV dataset and more densely sampled influenza data (1968 to 2003).

Topolow also effectively reduces the impact of experimental noise and bias –a critical feature for real-world applications where measurement errors are inevitable [11]. Overall, in all experiments, Topolow consistently outperformed MDS in terms of bias and variance of errors, outcome stability, noise reduction, and completeness of predicted antigenic distances on validation data.

Although we have focused on viral antigenic titers in this study, the Topolow algorithm can be used to characterize any continuous and relational phenotype of species that undergo directional selection pressure. The method’s ability to handle sparse, noisy data while maintaining biological relevance makes it particularly valuable for studying evolutionary trajectories in various biological systems, from pathogen adaptation to broader phenotypic evolution [42].

### 4.1 Future possibilities and limitations

Recent work has demonstrated the value of combining antigenic characterization with other data types for surveillance and vaccine strain selection [43, 44, 37, 45, 16, 17]. Topolow could enhance such efforts by providing comprehensive, accurate, and stable antigenic characterization across the viral strains. Topolow’s ability to predict antigenic phenotypes for under-characterized strains could be particularly valuable for early detection of emerging variants [46] and examining early indicators of cluster success [2, 47].

If there are islands in the graph of measurements in the data, Topolow can find the relational positions of the antigens in each island separately, but it is impossible to find the distances between members of distinct islands.

In initializing the positions by time, the evolution of antigens was assumed to be subject to a directional selection pressure, which holds true for viruses [21]. Otherwise, the temporal initialization should not be used, thus, the convergence of the positions may take longer.

The basic implementation of Topolow, as described in this paper, is time-intensive. There are various opportunities to improve it, such as a selective search scheme to skip moving the parts of the graph that have reached the acceptable proximity of their optimal location, use of C++, etc.

## 5 Authors’ statement

O.A. conceived the idea, developed the algorithm, and performed and analyzed the results. O.A. and P.R. designed the study framework and wrote the manuscript.

## 6 Acknowledgments

We gratefully acknowledge Derek Smith, Christian Gunning, Toby Brett, and Alpha Forna for valuable discussions and insights. This research was supported by federal funds from the National Institute of Allergy and Infectious Diseases, National Institutes of Health, Department of Health and Human Services, under Contract No. 75N93021C00018 (NIAID Centers of Excellence for Influenza Research and Response, CEIRR).

