## Supplementary Information for "Topolow: A mapping algorithm for antigenic cross-reactivity and binding affinity assay results"

### S0.1 The impact of missing values

Let $D$ be the matrix of distances of all test and reference antigens constructed from the available titers. For illustration, a $4 \times 4$ matrix may take the form:

$$\text{D} = \begin{array}{c} \\ \text{Ref. Ag 1} \\ \text{Ref. Ag 2} \\ \text{Test Ag 1} \\ \text{Test Ag 2} \end{array} \begin{array}{c} \begin{array}{cccc} \text{Ref. Ag 1} & \text{Ref. Ag 2} & \text{Test Ag 1} & \text{Test Ag 2} \end{array} \\ \begin{pmatrix} 0 & D_{12} & - & - \\ D_{21} & 0 & D_{23} & - \\ - & D_{32} & 0 & D_{34} \\ - & - & D_{43} & 0 \end{pmatrix} \end{array}$$

In this hypothetical example, there are 4 points, 3 measurements, and 6 missing entries. To identify the coordinates of the 4 points, MDS, specifically metric MDS, specifies an objective function $S(X)$:

$$S(X) = \sum_{i<j \& D_{ij} exists} \left(D_{ij} - \|x_i - x_j\|\right)^2 . \tag{S1}$$

where $x_i$ and $x_j$ are the locations of antigens $i$ and $j$ on the generated map. Typically, Gradient Descent or Newton's method is used to find the minimum for $S(X)$. Both algorithms are guided towards the solution by partial derivatives of the objective function. The partial derivative with respect to the $k$-th coordinate of point $i$ is:

$$\frac{\partial S}{\partial x_{ik}} = 2 \sum_{j \neq i \& D_{ij} exists} \left(\|x_i - x_j\| - D_{ij}\right) \frac{x_{ik} - x_{jk}}{\|x_i - x_j\|}. \tag{S2}$$

Notice that the formula for partial derivatives skips all the terms where $D_{ij}$ is missing from the summation. Therefore, the accuracy of magnitude and direction of the gradient vector $\frac{\partial \vec{S}}{\partial x_i}$ is affected by missing elements in $D$. On the other hand, Topolow performs pairwise distance optimizations sequentially for each pair of nodes, the optimization direction is simply the vector connecting them. By avoiding the need to compute a gradient vector, Topolow has the potential to operate robustly even when a substantial portion of entries in $D$ are missing.

### S0.2 Algorithm and implementation notes

We employ a cooling schedule where both spring stiffness $k$ and repulsion strength $c$ decrease according to a multiplicative function of iteration number. Specifically, at iteration $t$: $k(t) = k * (1 - \alpha)^t$ and $c(t) = c * (1 - \alpha)^t$ where $\alpha$ controls the cooling rate. This prevents oscillation and allows fine-scale adjustments in the final stages. The optimization terminates when the relative change in mean absolute error (MAE) between observed and projected antigenic distances falls below a preset threshold $\epsilon$: $|(MAE_t - MAE_{t-1})|/MAE_t < \epsilon$.

The **output** provides optimal $N$-dimensional coordinates for all test and reference antigens. While these coordinates can be used directly for distance calculations and downstream analyses, for visualization, Topolow projects high-dimensional coordinates into 2D using Principal Component Analysis (PCA), with appropriate scaling to preserve relative distances.

Due to random initialization, the absolute positions and orientations of mapped points may vary between runs. However, the relative distances and spatial relationships between points remain consistent across different executions of the algorithm, producing visually similar maps.

### S0.3 Likelihood analysis

The choice of error model plays a key role in antigenic mapping due to the heterogeneous nature of serological data. While it may be tempting to model HI titer measurement errors using a normal distribution given its common usage and mathematical convenience, a **Laplace** distribution appears more appropriate for this application [48]. HI titer measurements exhibit variability from multiple sources, including differences between laboratories, date of collection, observers, and assay details. Additionally, reporting low/high measurements as a threshold value introduces large errors. The Laplace distribution's **heavier tails** and greater robustness to outliers make it particularly well-suited for handling these heterogeneous sources of error. The Laplace model also aligns naturally with our **choice of using MAE** rather than squared error measures. The rationale for this choice is that larger errors often correspond to measurements at the limits of detection, and we do not want our optimization algorithm to be driven mostly by more inaccurate observations.

**Algorithm 1** TopoLow (Topological Optimization for Low-Dimensional Mapping)

**Require:**

$D$: $n \times n$ distance matrix
$k_0$: initial spring constant
$\alpha$: spring decay rate per iteration
$c$: repulsion constant
$N$: target dimensionality
$\epsilon$: convergence threshold :
$X$: $n \times N$ matrix of point coordinates in N-dimensional space :
$X \leftarrow n \times N$ ▷ matrix of initial coordinates
$k \leftarrow k_0$ ▷ Current springs constant
degrees $\leftarrow$ array of node degrees from $D$
**for** each pair of points $(i, j)$ where $i < j$ **do**
— $\delta \leftarrow X_j - X_i$ ▷ Vector between points
— $r \leftarrow \|\delta\|$ ▷ Current distance
— **if** $D_{i,j}$ is a measurement **then**
— — **if** $D_{i,j}$ is thresholded **then**
— — — **if** threshold condition not met **then**
— — — — $F \leftarrow (c/r^2)(\delta/r)$ ▷ Repulsive force
— — — **end if**
— — **else**
— — — $F \leftarrow k(D_{i,j} - r)(\delta/r)$ ▷ Spring force
— — **end if**
— — $X_i \leftarrow X_i - 2F/(4\text{degrees}_i + k)$
— — $X_j \leftarrow X_j + 2F/(4\text{degrees}_j + k)$
— **else**
— — $F \leftarrow (c/r^2)(\delta/r)$ ▷ Apply repulsive force for missing measurements
— — $X_i \leftarrow X_i - F/(2\text{degrees}_i)$
— — $X_j \leftarrow X_j + F/(2\text{degrees}_j)$
— **end if**
**end for**
Calculate MAE
**if** converged **then**
— break
**end if**
$k \leftarrow k(1 - \alpha)$ ▷ Decay spring constant
$c \leftarrow c(1 - \alpha)$ ▷ Decay repulsion constant

As mentioned, the distribution of errors is modeled by a **Laplace** model. The likelihood function under the Laplace model for errors of $n$ measurements, denoted as $e_i$, is:

$$L(\theta|e_1..e_n) = \prod_{i=1}^{n} \frac{1}{2b} \exp\left(-\frac{|e_i - \mu|}{b}\right), \tag{S3}$$

where $\mu$ and $b$ are location and scale parameters of the distribution. It can be shown that the maximum likelihood estimates (MLE) for $\mu$ and $b$ are $\hat{\mu} = \text{median}(e)$ and $\hat{b} = \frac{1}{n}\sum_{i=1}^{n} |e_i - \hat{\mu}|$. Since Laplace is a symmetric distribution, sample median can be estimated with sample mean: $\hat{\mu} = \text{mean}(e)$. Taking the negative log-likelihood and plugging in the MLEs:

$$\begin{aligned} NLL &= n\log(2b) + \frac{1}{b}\sum_{i=1}^{n} |e_i - \mu| \\ &= n\log(2b) + n = n\log(2\text{MAE}) + n. \end{aligned} \tag{S4}$$

#### S0.4 Adaptive Monte Carlo sampling for Parameter estimation

Unlike standard Monte Carlo methods that sample uniformly from the parameter space, the adaptive approach modifies the sampling distribution based on previously observed results. The **sampling probability is proportional** to the joint distribution of NLL. Thus, to obtain each sample, we reconstruct the NLL surface in the space of all parameters using a kernel density estimator (KDE). This approach concentrates sampling in regions of the parameter space that are more likely to contain optimal values.

To evaluate the **validation error** with each set of parameters, we used **k-fold cross-validation** with $k = 20$. For each fold, we randomly removed 5% of measurements for testing while maintaining the remaining data.

To initialize the **parameter search**, we generate 50 parameter combinations using **Latin** hypercube sampling [49], ensuring broad and efficient coverage of the parameter space. The algorithm then draws additional samples (typically 1000), updating the sampling distribution after each iteration. The peak of the final **likelihood surface** determines the optimal parameter values. Fig. S-4 and S-5 show profile likelihoods and 95% marginal confidence intervals for all parameters with H3N2 and HIV data sets.

##### S0.4.1 Visual inspection

Figure S-3 shows the maps created by Topolow and MDS for our most challenging scenario (10 dimensions, 95% missing distances, with added noise and bias). This high quality of preservation of global structure is particularly noteworthy given the extreme sparsity and noise in this scenario.

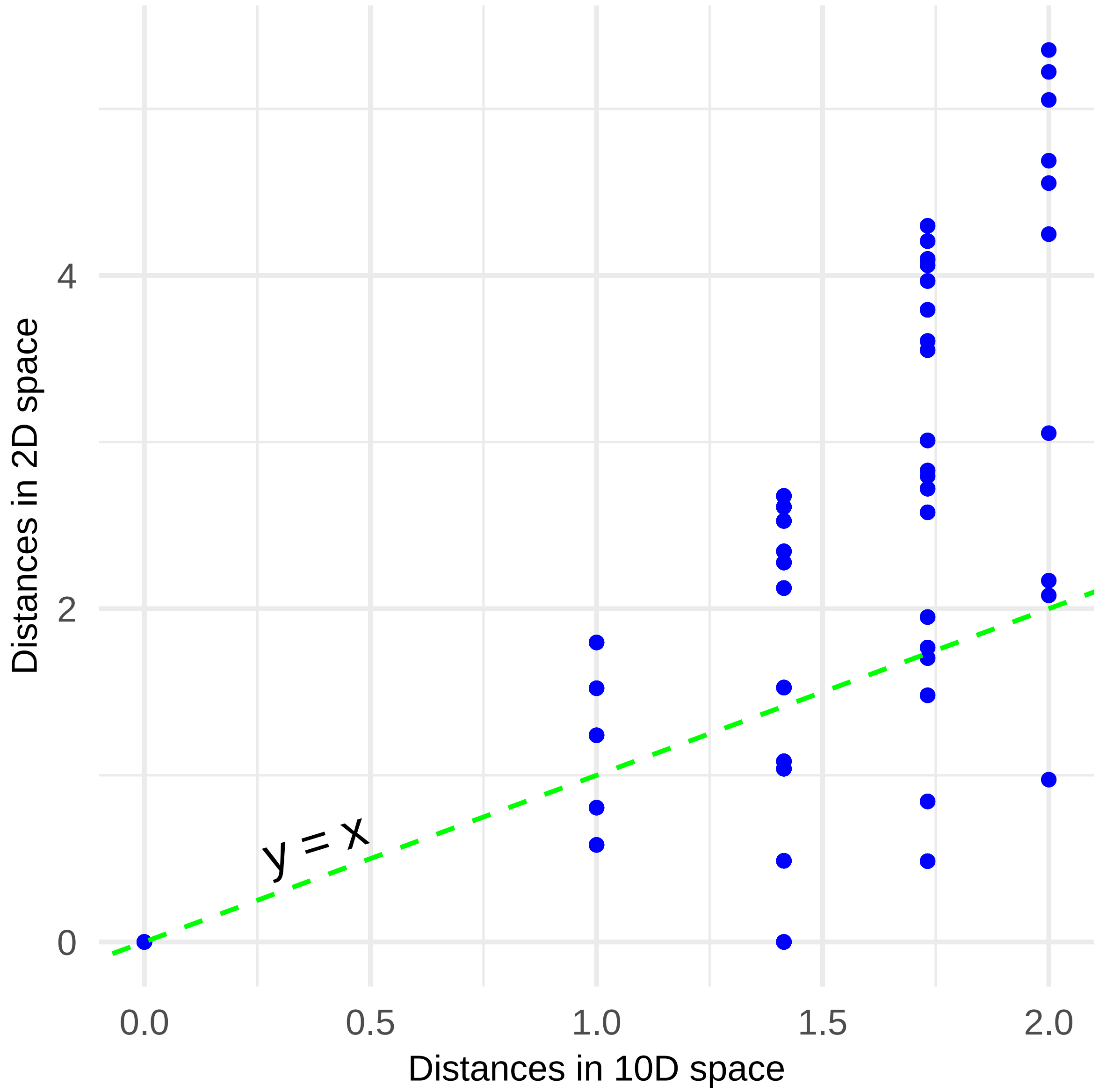


Figure S-1: The pairwise distances in the original space (10D) versus the pairwise distances after reducing the dimensions to 2 using the MDS method applied to 15 points spread uniformly in a 10D space.

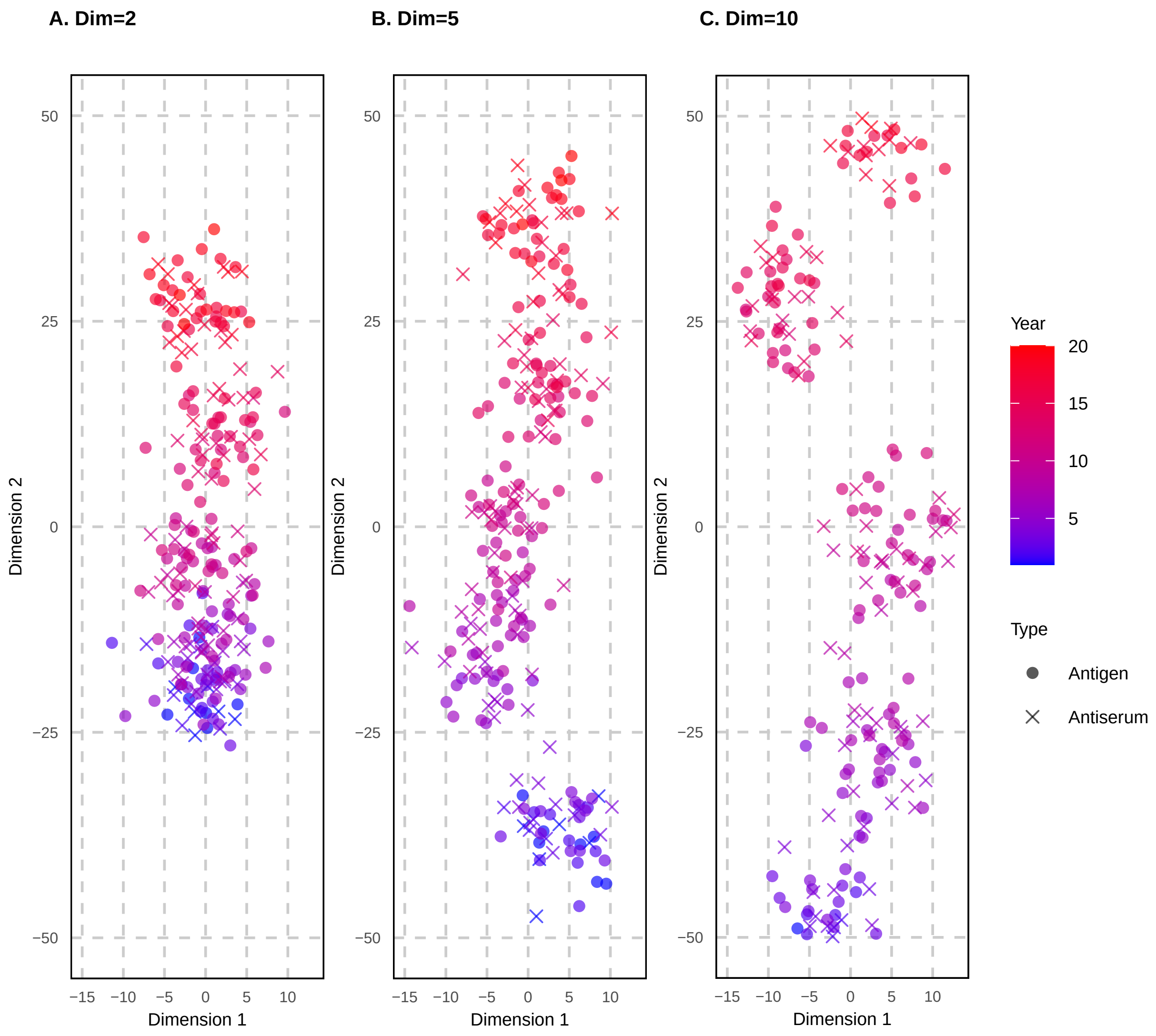


Figure S-2: 2D projection of three simulated datasets using PCA. Imaginary years 1 to 20 are assigned to points. Note that the difference in trend strengths and variation in dispersity observed in 2D is a byproduct of the differences in original dimensionalities of datasets.

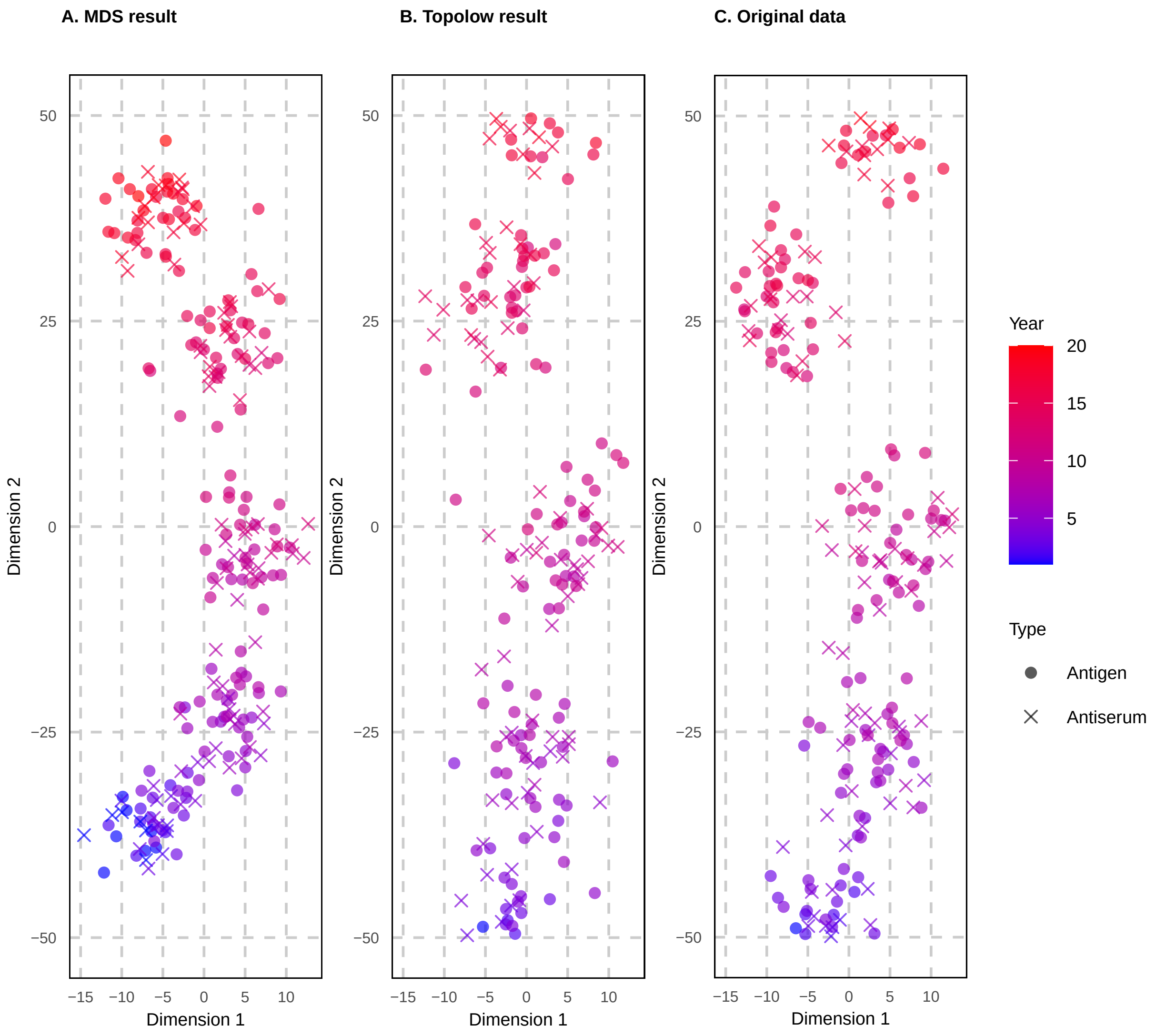


Figure S-3: Antigenic maps for simulated scenario with 10 dimensions, 95% missing distances, and added noise and bias using MDS (**A**) and Topolow (**B**) compared to the 2D projection of the original data (**C**).

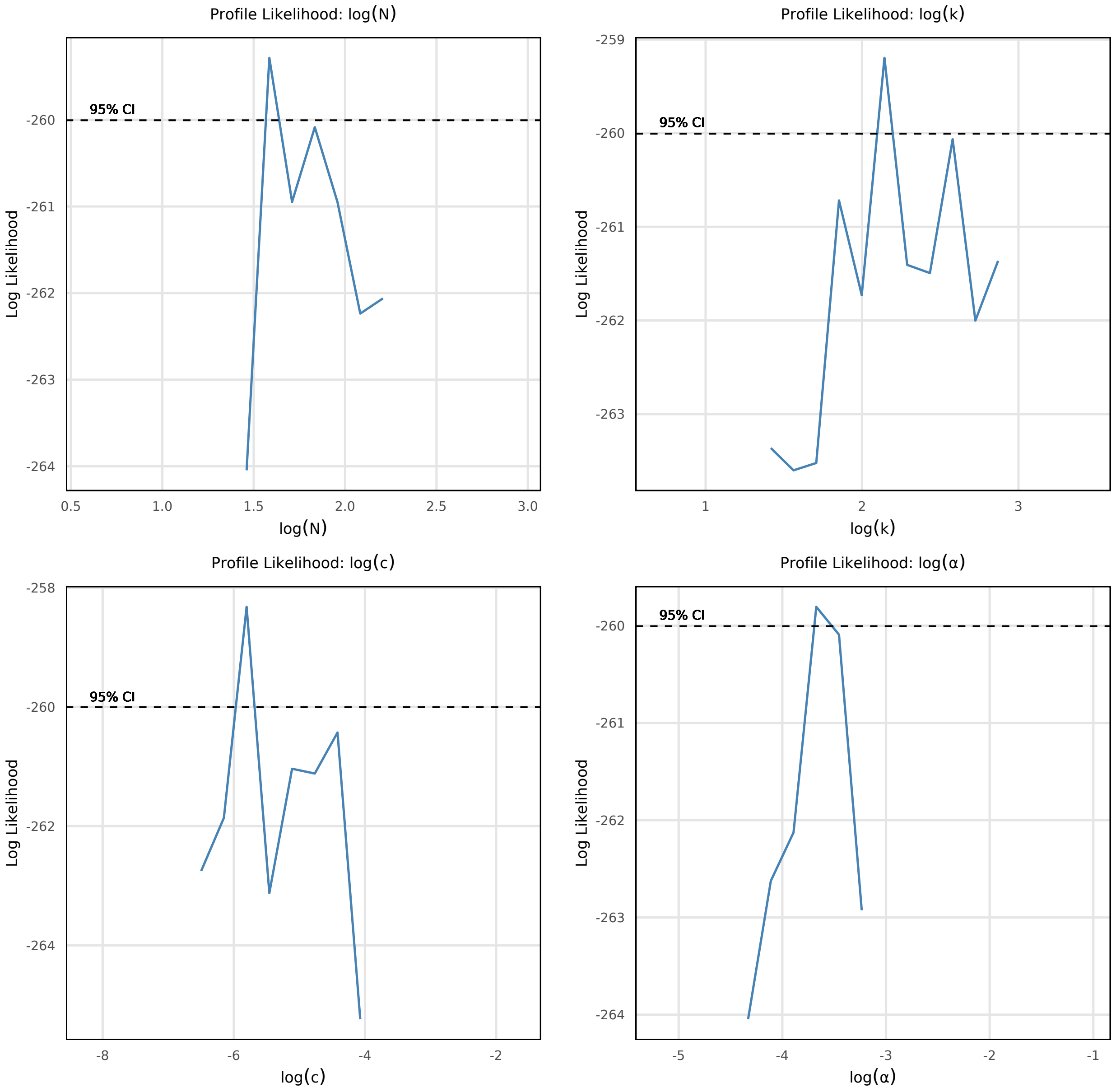


Figure S-4: Profile likelihood plots for model parameters with H3N2 dataset. Profile likelihood for (a) dimensionality parameter $N$, (b) initial spring constant $k$, (c) repulsion constant $c$, and (d) cooling rate $\alpha$, showing optimal value and 95% confidence interval.

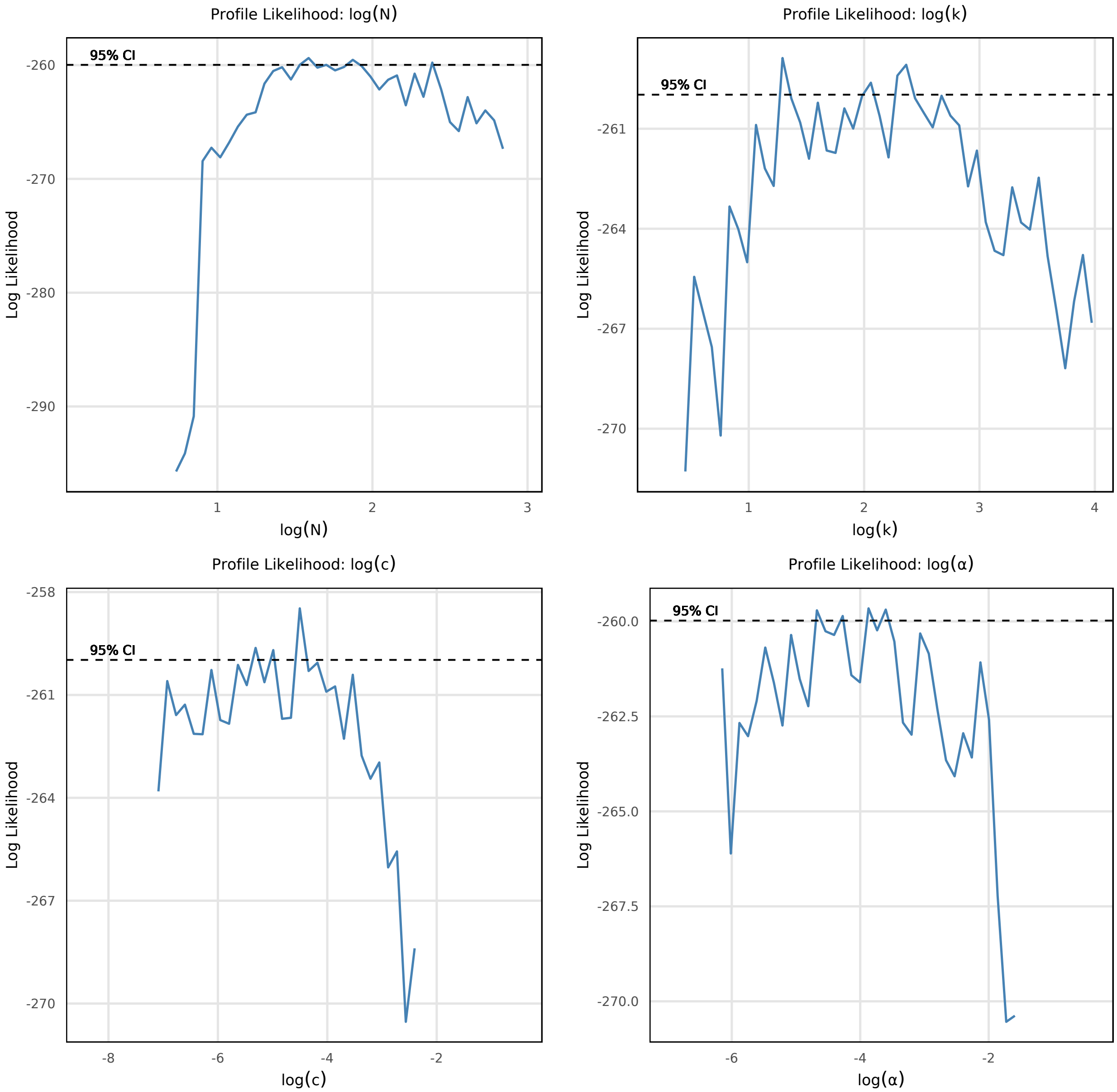


Figure S-5: Profile likelihood plots for model parameters with HIV dataset. Profile likelihood for (a) dimensionality parameter $N$, (b) initial spring constant $k$, (c) repulsion constant $c$, and (d) cooling rate $\alpha$, showing optimal value and 95% confidence interval.

Table S-1: Count of mapped antigens with simulated data. For each scenario, MDS was ran with 2 dimensions and the same number of dimensions as Topolow (the optimal dimensionality found by Topolow.)

| Dim | Missing | MDS (2D) | MDS (Opt. Dim.) | Topolow |
|---|---|---|---|---|
| 2 | 70% | 250 | 250 | 250 |
| 2 | 85% | 250 | 250 | 250 |
| 2 | 95% | **246** | **240** | 250 |
| 5 | 70% | 250 | 250 | 250 |
| 5 | 85% | 250 | 250 | 250 |
| 5 | 95% | **248** | **30** | 250 |
| 10 | 70% | 250 | **188** | 250 |
| 10 | 85% | 250 | **112** | 250 |
| 10 | 95% | **245** | **3** | 250 |

Table S-2: Comparison of MAPE (%) in the inputs due to added noise/biase against MAPE of locations determined by MDS and Topolow.

| Dim. | Missing | Variant | Input | MDS | Topolow |
|---|---|---|---|---|---|
| 2 | 70% | +Noise | 8.046 | 12.630 | 3.764 |
| 2 | 70% | +Noise+Bias | 11.121 | 12.723 | 5.895 |
| 2 | 85% | +Noise | 7.888 | 19.133 | 4.496 |
| 2 | 85% | +Noise+Bias | 11.133 | 18.821 | 6.243 |
| 2 | 95% | +Noise | 7.141 | 30.696 | 5.872 |
| 2 | 95% | +Noise+Bias | 10.183 | 29.355 | 7.162 |
| 5 | 70% | +Noise | 5.693 | 29.917 | 3.205 |
| 5 | 70% | +Noise+Bias | 8.488 | 29.769 | 6.507 |
| 5 | 85% | +Noise | 5.420 | 37.041 | 4.073 |
| 5 | 85% | +Noise+Bias | 7.867 | 35.523 | 6.328 |
| 5 | 95% | +Noise | 4.791 | 48.159 | 5.459 |
| 5 | 95% | +Noise+Bias | 7.173 | 47.703 | 7.324 |
| 10 | 70% | +Noise | 5.411 | 37.265 | 3.825 |
| 10 | 70% | +Noise+Bias | 7.989 | 37.121 | 6.504 |
| 10 | 85% | +Noise | 5.170 | 44.970 | 4.559 |
| 10 | 85% | +Noise+Bias | 7.507 | 45.315 | 6.634 |
| 10 | 95% | +Noise | 4.855 | 53.920 | 5.343 |
| 10 | 95% | +Noise+Bias | 6.900 | 53.654 | 6.910 |

Table S-3: Paired t-test results between validation MAE of Topolow and MDS over each fold of a 20-fold cross-validation simulation for all scenarios.

| Dim. | Missing | Variant | p-value |
|---|---|---|---|
| 2 | 70% | +Noise | 0e+00 |
| 2 | 70% | +Noise+Bias | 0e+00 |
| 2 | 70% | Original | 0e+00 |
| 2 | 85% | +Noise | 0e+00 |
| 2 | 85% | +Noise+Bias | 0e+00 |
| 2 | 85% | Original | 0e+00 |
| 2 | 95% | +Noise | 1e-07 |
| 2 | 95% | +Noise+Bias | 0e+00 |
| 2 | 95% | Original | 0e+00 |
| 5 | 70% | +Noise | 0e+00 |
| 5 | 70% | +Noise+Bias | 0e+00 |
| 5 | 70% | Original | 0e+00 |
| 5 | 85% | +Noise | 0e+00 |
| 5 | 85% | +Noise+Bias | 0e+00 |
| 5 | 85% | Original | 0e+00 |
| 5 | 95% | +Noise | 0e+00 |
| 5 | 95% | +Noise+Bias | 0e+00 |
| 5 | 95% | Original | 0e+00 |
| 10 | 70% | +Noise | 0e+00 |
| 10 | 70% | +Noise+Bias | 0e+00 |
| 10 | 70% | Original | 0e+00 |
| 10 | 85% | +Noise | 0e+00 |
| 10 | 85% | +Noise+Bias | 0e+00 |
| 10 | 85% | Original | 0e+00 |
| 10 | 95% | +Noise | 0e+00 |
| 10 | 95% | +Noise+Bias | 0e+00 |
| 10 | 95% | Original | 0e+00 |

Table S-4: Paired t-tests between errors of Topolow and MDS in empirical datasets.

| Dataset | Dimension | Missing | p-value |
|---|---|---|---|
| Influenza (H3N2) | 2 | 91% | 0e+00 |
| HIV (B & C) | 2 | 94% | 0e+00 |

Table S-5: Performance across complexity levels and missingness proportions (MAPE ± SD). Supporting information for Figure 2

| Dim. | Missing | Variant | MDS | Topolow |
|---|---|---|---|---|
| 2 | 70% | Original | 13.261 ± 0.668 | 0.536 ± 0.975 |
| 2 | 70% | +Noise | 12.435 ± 0.835 | 4.650 ± 0.982 |
| 2 | 70% | +Noise+Bias | 12.753 ± 0.899 | 5.860 ± 0.383 |
| 2 | 85% | Original | 18.575 ± 1.620 | 0.046 ± 0.037 |
| 2 | 85% | +Noise | 19.057 ± 2.591 | 6.103 ± 1.825 |
| 2 | 85% | +Noise+Bias | 18.929 ± 1.980 | 6.447 ± 0.947 |
| 2 | 95% | Original | 32.629 ± 5.457 | 5.608 ± 4.023 |
| 2 | 95% | +Noise | 32.028 ± 4.812 | 16.766 ± 6.073 |
| 2 | 95% | +Noise+Bias | 31.772 ± 5.895 | 13.975 ± 5.032 |
| 5 | 70% | Original | 31.462 ± 1.074 | 0.385 ± 0.688 |
| 5 | 70% | +Noise | 29.875 ± 1.519 | 4.071 ± 0.465 |
| 5 | 70% | +Noise+Bias | 29.699 ± 1.093 | 6.672 ± 0.449 |
| 5 | 85% | Original | 37.204 ± 1.557 | 0.162 ± 0.084 |
| 5 | 85% | +Noise | 36.671 ± 1.446 | 6.816 ± 0.999 |
| 5 | 85% | +Noise+Bias | 35.188 ± 2.037 | 7.266 ± 0.662 |
| 5 | 95% | Original | 45.063 ± 3.037 | 19.186 ± 5.565 |
| 5 | 95% | +Noise | 44.151 ± 2.785 | 19.886 ± 3.908 |
| 5 | 95% | +Noise+Bias | 44.227 ± 2.873 | 18.197 ± 4.934 |
| 10 | 70% | Original | 39.750 ± 1.498 | 0.083 ± 0.023 |
| 10 | 70% | +Noise | 37.252 ± 1.587 | 5.831 ± 0.524 |
| 10 | 70% | +Noise+Bias | 37.174 ± 1.451 | 7.198 ± 0.458 |
| 10 | 85% | Original | 46.636 ± 1.879 | 2.491 ± 0.866 |
| 10 | 85% | +Noise | 44.639 ± 1.921 | 8.423 ± 0.759 |
| 10 | 85% | +Noise+Bias | 44.728 ± 1.755 | 8.166 ± 0.678 |
| 10 | 95% | Original | 52.486 ± 2.820 | 15.425 ± 2.915 |
| 10 | 95% | +Noise | 50.271 ± 2.378 | 15.784 ± 2.997 |
| 10 | 95% | +Noise+Bias | 50.753 ± 2.899 | 14.389 ± 2.478 |

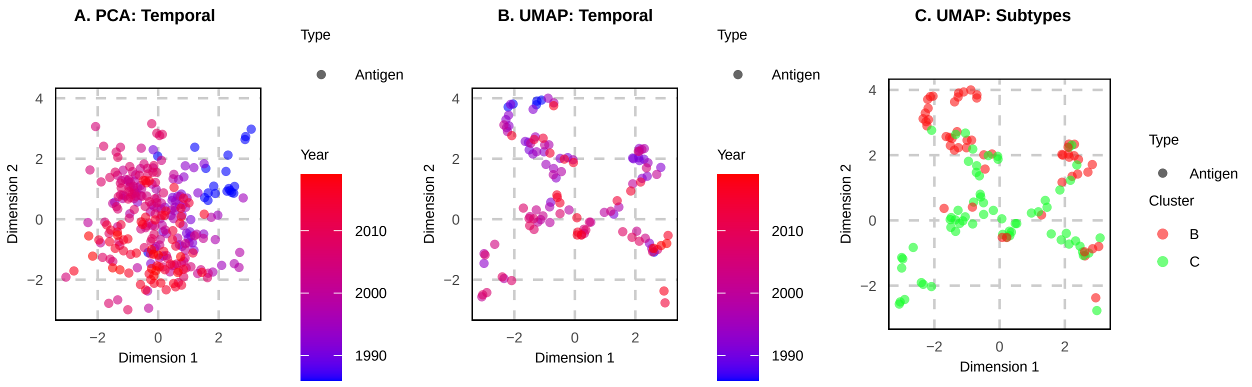


Figure S-6: 2D visualization of HIV antigenic map. Panel (**A**) shows the temporal progression of antigenic change, with smaller diversity in the older isolates and larger diversity in more recent ones; (**B**) and (**C**) show the UMAP transformation of the same map. (C) exposes the clusters in the data, based on the two subtypes, more clearly.

Table S-6: Distribution of errors (Mean ± SD). Mean values quantify the bias in estimates by each method.

| Dim. | Missing | Variant | MDS | Topolow |
|---|---|---|---|---|
| 2 | 70% | +Noise | 1.089 ± 1.028 | -0.059 ± 0.308 |
| 2 | 70% | +Noise+Bias | 1.203 ± 0.962 | 0.533 ± 0.394 |
| 2 | 70% | Original | 1.307 ± 1.017 | 0.014 ± 0.302 |
| 2 | 85% | +Noise | 1.377 ± 1.345 | -0.059 ± 0.359 |
| 2 | 85% | +Noise+Bias | 1.462 ± 1.343 | 0.427 ± 0.409 |
| 2 | 85% | Original | 1.500 ± 1.317 | -0.001 ± 0.005 |
| 2 | 95% | +Noise | 1.805 ± 2.177 | -0.213 ± 1.318 |
| 2 | 95% | +Noise+Bias | 1.731 ± 2.181 | 0.153 ± 1.217 |
| 2 | 95% | Original | 1.868 ± 2.146 | -0.116 ± 1.015 |
| 5 | 70% | +Noise | 5.103 ± 2.709 | -0.067 ± 0.727 |
| 5 | 70% | +Noise+Bias | 5.070 ± 2.657 | 1.021 ± 0.814 |
| 5 | 70% | Original | 5.424 ± 2.584 | 0.005 ± 0.269 |
| 5 | 85% | +Noise | 5.746 ± 2.993 | -0.161 ± 1.199 |
| 5 | 85% | +Noise+Bias | 5.523 ± 2.932 | 0.833 ± 1.058 |
| 5 | 85% | Original | 5.869 ± 2.912 | -0.005 ± 0.096 |
| 5 | 95% | +Noise | 6.070 ± 3.862 | -0.837 ± 2.959 |
| 5 | 95% | +Noise+Bias | 6.066 ± 3.826 | -0.191 ± 3.344 |
| 5 | 95% | Original | 6.127 ± 3.806 | -0.791 ± 3.572 |
| 10 | 70% | +Noise | 9.202 ± 3.038 | -0.320 ± 1.497 |
| 10 | 70% | +Noise+Bias | 9.229 ± 3.055 | 1.346 ± 1.574 |
| 10 | 70% | Original | 9.893 ± 3.013 | -0.003 ± 0.049 |
| 10 | 85% | +Noise | 9.994 ± 3.345 | -0.496 ± 1.998 |
| 10 | 85% | +Noise+Bias | 10.023 ± 3.412 | 0.961 ± 1.925 |
| 10 | 85% | Original | 10.520 ± 3.284 | -0.050 ± 0.858 |
| 10 | 95% | +Noise | 10.003 ± 4.546 | -0.999 ± 4.028 |
| 10 | 95% | +Noise+Bias | 9.921 ± 4.590 | 0.248 ± 3.780 |
| 10 | 95% | Original | 10.289 ± 4.446 | -0.848 ± 4.128 |
